# Metabolic implications for dual substrate growth in “*Candidatus* Accumulibacter”

**DOI:** 10.1101/2025.01.13.628880

**Authors:** Timothy Páez-Watson, Casper Jansens, Mark C.M. van Loosdrecht, Samarpita Roy

## Abstract

This study explores the metabolic implications of dual substrate uptake in *“Candidatus Accumulibacter”*, focusing on the co-consumption of volatile fatty acids and amino acids under conditions typical of enhanced biological phosphorus removal (EBPR) systems. Combining batch tests from highly enriched “*Ca.* Accumulibacter” cultures with conditional flux balance analysis (cFBA) predictions, we demonstrated that co-consumption of acetate and aspartate leads to synergistic metabolic interactions, lowering ATP loss compared to individual substrate consumption. The metabolic synergy arises from the complementary roles of acetate and aspartate uptake: acetate uptake provides acetyl-CoA to support aspartate metabolism, while aspartate conversion generates NADH, reducing the need for glycogen degradation during acetate uptake. We termed this type of metabolic interaction as reciprocal synergy. We further expanded our predictions to uncover three types of interactions between catabolic pathways when substrates are co-consumed by “*Ca.* Accumulibacter”: (i) neutral, (ii) one-way synergistic and (iii) reciprocal synergistic interactions. Our results highlight the importance of network topology in determining metabolic interactions and optimizing resource use. These findings provide new insights into the metabolism “*Ca.* Accumulibacter” and suggest strategies for improving EBPR performance in wastewater treatment plants, where the influent typically contains a mixture of organic carbon compounds.

**Synopsis:** This research demonstrates how dual substrate uptake by “*Ca.* Accumulibacter” enhances metabolic efficiency in EBPR by reducing global ATP losses through optimization of storage polymer usage.

## 1. Introduction

Enhanced biological phosphorous removal (EBPR) systems are complex, dynamic environments that foster the growth of mixed microbial communities. Amongst these, members of the genus “*Ca.* Accumulibacter” are dominant, both in terms of biovolume^1, 2^ and protein content^3, 4^, highlighting their key role in the biological conversions that drive this process. Their unique metabolic capabilities allow them to accumulate large amounts of polyphosphate (PolyP)^5, 6^ under cyclic, dynamic environments. No pure cultures of “*Ca.* Accumulibacter” are available, making research reliant on highly enriched cultures (> 90 % biovolume^1, 2^) for characterization.

The metabolism of “*Ca.* Accumulibacter” depends on the cycling of polyhydroxyalkanoates (PHAs), PolyP and glycogen during an EBPR cycle^7–9^. This enables the rapid anaerobic uptake and accumulation of volatile fatty acids (VFAs) inside the microbial cells. The stored polymers can be subsequently oxidized aerobically, generating enough energy for growth and the accumulation of PolyP. Research on this metabolic strategy has predominantly focused on feeding single substrates, such as VFAs like acetate ^10–12^ and propionate ^13^, as well as non-VFA substrates like glucose ^14, 15^, and amino acids like aspartate and glutamate^16^.

It is noteworthy that environments containing only a single substrate are rare in nature, and wastewater treatment plants are no exception. Wastewater typically contains a mixture of organic substrates, including fatty acids, amino acids, sugars and lipids^17–19^. Understanding the effects of multiple substrates on the metabolism of microorganisms involved in phosphate removal is therefore crucial for optimizing EBPR processes.

Despite the importance of mixed substrates, studies on the metabolic mechanisms of “*Ca.* Accumulibacter” during the simultaneous uptake of two or more substrates are very limited. Studies have reported co-consumption of acetate with glucose ^15, 20^, acetate with propionate and lactate ^21^ and acetate with glycerol^22^ in enrichment cultures, but the interactions between the metabolic strategies for consumption of each substrate remains poorly understood.

A notable study by Qiu et al. ^16^ stands out for its detailed examination of the concurrent uptake of acetate with either aspartate or glutamate. This research identified a potential synergy between acetate and aspartate, where aspartate uptake led to a net energy gain (∼9 % ATP gain), enhancing acetate uptake. The proposed mechanism was the operation of fumarate reductase (reducing fumarate to succinate) which contributes to the *proton motive force* (pmf) required for acetate transport. More recently, a similar energetic gain was evaluated for acetate and succinate co-consumption^23^, though the entry point of succinate bypasses fumarate reductase, raising the question of whether this synergy is indeed driven by *pmf* or an unidentified metabolic interaction.

Understanding the interactions within metabolic networks is challenging due to the complexity and interconnectedness of metabolites, especially energy carriers like ATP, NADH, FADH_2_^24^. In this regard, metabolic modelling provides valuable tools for studying these networks and how interactions emerge from stoichiometric rules. Techniques such as Flux Balance Analysis (FBA) can model steady-state metabolic operations^25^, but more advanced methods are needed for dynamic systems like EBPR^26–28^. One such method is conditional FBA (cFBA)^29^, which has been successfully used to model “*Ca.* Accumulibacter” metabolism, where intracellular storage polymers cycling emerged as a property of model stoichiometry and environmental conditions^30^. However, how these emergent properties change with multiple substrates remains an open question.

To address this, we aimed to understand the metabolic implications of dual substrate uptake in “*Ca.* Accumulibacter” under the dynamic conditions characteristic of EBPR. We developed and tested a metabolic model for the uptake of varying acetate and aspartate ratios to uncover synergistic interactions that improve growth yields. These findings were validated with lab-scale enrichments and the mechanisms behind the synergy are described. Finally, we extended this modelling approach to explore interactions with additional substrates, identifying a basic biological principle of metabolic interactions. This work lays the groundwork for further exploration of multiple substrate consumption, which could potentially lead to increased biomass yields compared to individual substrate consumption.

## 2. Materials and Methods

### Metabolic model and cFBA simulations

The metabolism of “*Ca.* Accumulibacter” was simulated with cFBA using the py_cFBA toolkit implementation^31^. A basic metabolic model was constructed as a stoichiometric matrix (**S**), representing the relationships between metabolites and reactions. Stoichiometries for reactions involved in glycogen degradation, glycolysis, the TCA cycle, anaplerotic routes, and PHA synthesis were adapted from an earlier study on “*Ca.* Accumulibacter”^32^, excluding the reaction *MalE* which was present in only a few genomes within this genus. Stoichiometries for aspartate metabolism were obtained from^16^ and the presence of this pathway in our enrichment culture was confirmed with metagenomics (see later in this section). A reaction representing synthesis of 1 c-mole of biomass was implemented in **S** following the stoichiometry from ^30^, which combined the energy (ATP) requirements for bacterial growth from acetyl-CoA from ^33^ and the overall stoichiometry of PAOs growth from ^8, 34^.

In the model, selected metabolites were defined as imbalanced, allowing their accumulation or depletion over time during simulations. These included acetate, aspartate, glycogen, PHB, PH2MV, CO_2_, polyP, and biomass. All other metabolites were balanced, adhering to the steady-state assumption of FBA. Biomass was defined as the sole contributor to the weights vector (*w*) in the cFBA formulation. The **S** matrix, together with details on imbalanced metabolites and the weights vector, is provided in the supplementary materials.

The model was implemented in Python using the py_cFBA toolkit, which generated SBML files for each configuration. Simulations of an EBPR cycle consisted of five time points (Δt = 1 hour), with no enzyme capacity constraints. Anaerobic and aerobic phases were simulated by allowing the reactions ETC_NADH, ETC_FADH (electron transport chain oxygen consumption), and biomass synthesis to occur exclusively in the final two time points of each cycle. Reaction reversibility was defined using upper and lower bounds based on a prior thermodynamic evaluation^32^ (see supplementary information for each reaction definition).

Substrate uptake during the anaerobic phase was enforced using quota definitions. An equality-quota at the initial time point specified the concentration of substrate fed, followed by a max-quota of zero in subsequent time points. This enforced the anaerobic uptake of substrate. Substrate concentrations were normalized to provide equivalent electron equivalents, even for substrate mixtures, based on their degree of reduction (e.g., 8 electrons for acetate, 12 electrons for aspartate). All simulations optimized biomass synthesis as the global target across the entire cycle, rather than at each time step, consistent with cFBA methodology. All SBML models, simulation files and results are available at https://github.com/TP-Watson/PAOs_co-substrates_cFBA.

### “*Ca* Accumulibacter” Enrichment

A “Ca. Accumulibacter” enrichment was obtained in a 1.5 L (1 L working volume) sequencing batch reactor (SBR), following conditions described earlier^2^ with some adaptations. The reactor was inoculated using enriched sludge from the work of Páez-Watson et al.^2^, which was previously inoculated with activated sludge from a municipal wastewater treatment plant (Harnaschpolder, The Netherlands). Each SBR cycle lasted 6 hours, consisting of 30 minutes of settling, 50 minutes of effluent removal, 10 minutes of N_2_ sparging, 30 minutes of anaerobic feeding, 105 minutes of anaerobic phase and 135 minutes of aerobic phase. N_2_ gas and compressed air were sparged at 500 ml/min into the reactor broth to maintain anaerobic and aerobic conditions respectively. The hydraulic retention time (HRT) was 12 hours (removal of 500 mL of broth per cycle, each cycle of 6 hours). The average solids retention time (SRT) was controlled to 9 days by the removal of 27,7 ml of mixed broth at the end of the mixed aerobic phase in each cycle. The pH was controlled at 7.3 ± 0.1 by dosing 0.5 M HCl or 0.5 M NaOH. The temperature was maintained at 20 ± 1 °C.

The reactor was fed with three separate media components diluted in demineralized water: a concentrated COD medium (400 mg COD/L) of acetate (13 g/L sodium acetate ×3H_2_O); a concentrated mineral medium (0.69 g/L NH_4_Cl, 2.16 g/L MgSO_4_×7H_2_O, 0.54 g/L CaCl_2_×2H_2_O, 0.64 KCl, 0.06 g/L N-allylthiourea (ATU), 0.06 g/L yeast extract and 6 mL/L of trace element solution prepared following Smolders et al.^7^; and a phosphate solution containing 0.76 g/L NaH_2_PO_4_×H_2_O and 0.8 g/L Na_2_HPO_4_×2H_2_O. In each cycle, 75 mL of COD medium, 75 ml of mineral medium and 360 mL of phosphate solution were added to the reactor during the 30 minutes of feeding. The final feed contained 400 mg COD/L of acetate.

### Batch tests

Batch tests were conducted in the bioreactor on the enriched biomass once a *pseudo* steady state was reached (determined by a constant phosphate release and removal over multiple days). For the batch tests, 400 ml of H_2_O and 50 ml of mineral media were fed as usual during the anaerobic phase. Later, 50 ml of organic substrate (containing either acetate, aspartate or a mix) was pulse fed and considered the beginning of the anaerobic phase of the cycle. The anaerobic phase on these batch tests was extended by 30 minutes to compensate for the delay in feed. The organic media was prepared such that the final feed contained 400 mg COD/L of acetate, aspartate or a mix (calculated by using the degree of reduction of 8 and 12 e^-^/mol for acetate and aspartate, respectively). Thus, the organic substrate solution for the tests contained only acetate, only aspartate or mix of acetate and aspartate as follows: (i) 13.1 g/L sodium acetate trihydrate (C₂H₃NaO₂·3H₂O), (ii) 19.35 g/L sodium aspartate (C₄H₆NNaO₄), or (iii) 5,8 g/L sodium acetate trihydrate with 9,6 g/L sodium aspartate. For mixed substrates, the net consumption of acetate and aspartate was used to determine the acetate:aspartate uptake ratio.

### Reactor and biomass analyses

Extracellular concentrations of phosphate and ammonium were measured with a Gallery Discrete Analyzer (Thermo Fisher Scientific, Waltham, MA). Acetate was measured by high performance liquid chromatography (HPLC) with an Aminex HPX-87H column (Bio-Rad, Hercules, CA), coupled to RI and UV detectors (Waters, Milford, MA), using 0.0015 M phosphoric acid as eluent supplied at a flowrate of 1 mL/min.

The biomass concentration (total and volatile suspended solids – TSS and VSS) was measured in accordance with Standard Methods as described in Smolders et al.^7^ with some modifications: 10 ml of mixed broth were obtained at the end of the aerobic phase, centrifuged at 3600*g during 3 minutes and washed twice with demineralized water to remove salts. The sludge was then dried at 100 °C for 24 hours and weighed on a microbalance to determine the dry content – TSS. The ash content was determined by incinerating the dry material in an oven at 550 °C, and the difference used to calculate the VSS.

For glycogen and PHA determination, biomass samples (10 ml mixed broth) were collected throughout the batch test and stored in 15 ml conical tubes containing 0.3 ml of 37 % formaldehyde to stop biological activity. After each batch test, the biomass tubes were pottered to break the granular structure of the biomass, centrifuged at 3700 *g for 5 minutes and washed twice. The pellet was then frozen at −80 °C for at least 3 hours and freeze dried. For glycogen analysis the method described by Smolders et al.^7^ was used: 5 mg of dry biomass was digested in 0.9 M HCl solutions in glass tubes at 100 °C for 5 hours. After this time, tubes were cooled at room temperature, filtered with 0.45 µm Whatman disk filters and neutralized with equal volumes of 0.9 M NaOH. The glucose resulting from digestion was quantified using the D-Glucose Assay Kit (GOPOD Format) from Megazyme (Bray, Ireland).

### Microbial community characterization

The microbial community of the reactor was characterized at pseudo steady state as defined earlier. Two orthogonal approaches were used for the community characterization: metagenomics and Fluorescence in-situ hybridization (FISH).

For FISH, samples underwent handling, fixation, and staining procedures outlined by Winkler et al^35^. Bacteria were selectively identified using a blend of EUB338, EUB338-II, and EUB338-III probes^36, 37^. “*Ca*. Accumulibacter” was visualized employing mixtures of the probes Acc1011, Acc471, Acc471_2, Acc635, Acc470 designed and tested previously for different “*Ca.* Accumulibacter” lineages^38^. The images were captured with an epifluorescence microscope equipped with filter set Cy3 (ET545/25x ET605/70 m T565LPXR), Cy5 (ET640/30x ET690/50 m T660LPXR), and FITC (ET470/40x ET525/50 m T495LPXR) (Axio Imager M2, Zeiss, Germany). Quantitative FISH (qFISH) was done as a percentage of total biovolume over 12 representative pictures using the Daime software (DOME, Vienna, Austria)^39^.

For metagenomics, DNA from the biomass samples was extracted using the DNeasy PowerSoil Pro-Kit (Qiagen, Germany) following the manufacturer’s protocol. Shotgun sequencing was performed by Hologenomix (Delft, Netherlands). Paired-end sequencing with a read length of 150 bp was conducted using the Illumina NovaSeq X sequencing system. Library preparation was carried out using the Nextera XT DNA Library Preparation Kit. Approximately 10 Gbp of sequencing data were generated per sample.

The quality of raw sequenced reads was assessed using FastQC (version 0.11.7) with default parameters^40^, and results were visualized with MultiQC (version 1.19). Low-quality paired-end reads were trimmed and filtered using Fastp (version 0.23.4) in paired-end mode^41^. Taxonomic classification of raw reads was performed to profile the microbiome in each sample using Kraken2 (version 2) with the standard database, which includes all complete bacterial, archaeal, and viral genomes in the NCBI RefSeq database, complemented by a curated wastewater database (sludgeDB)^42^.

Clean reads were assembled into contigs using MetaSPAdes (version 3.15.5) with default parameters^43^. The resulting contigs were binned using MetaBAT (version 2.2.15) to reconstruct metagenome-assembled genomes (MAGs) with default parameters^44^. Bin completeness and contamination were assessed using CheckM (version 1.2.2) with the “lineage_wf” workflow ^45^. Relative abundance of bins with contamination below 5% was determined in each sample using CoverM (version 0.7.0, https://github.com/wwood/CoverM) with default parameters.

For phylogenetic analysis, bins were classified using GTDB-Tk (version 2.4.0) and GTDB release 220^46^. The *ppk1* gene was utilised as a marker in bins identified as Accumulibacter. hmmsearch ^47^ was used with the ppk1.hmm profile, taking the best hit as the *ppk1* gene. Identified *ppk1* genes were combined with those in an existing database and aligned with MUSCLE (version 5.1)^48^. A phylogenetic tree of these *ppk1* sequences was generated with RaxML-NG (version 1.2.2)^49^ and visualised using iTol (v6)^50^. Sequencing data was deposited under the BioProject no. PRJNA1191880, BioSample accession no. SAMN45106973.

## 3. Results

### Different anaerobic stoichiometries are employed for acetate, aspartate and combined substrate uptake by “*Ca.* Accumulibacter”

We expanded a previous metabolic model of “*Ca.* Accumulibacter”^32^ to incorporate uptake mechanisms for acetate and aspartate (Figure 1.A). Next, we simulated the concurrent uptake of both substrates at varying ratios. To normalize the simulations, we ensured that the combined substrates provided the same amount of electron equivalents (analogous to the chemical oxygen demand, most widely used in engineering), meaning the ratios were adjusted to achieve ‘electron equivalence’ rather than molar equivalence (see methods for details). The predictions indicated that the optimal anaerobic strategies employed for the individual uptake of acetate or aspartate were different, and that a mixed uptake also resulted in different individual anaerobic strategies rather than linear combinations of the individual strategies (Figure 1.B).

**Figure 1.**
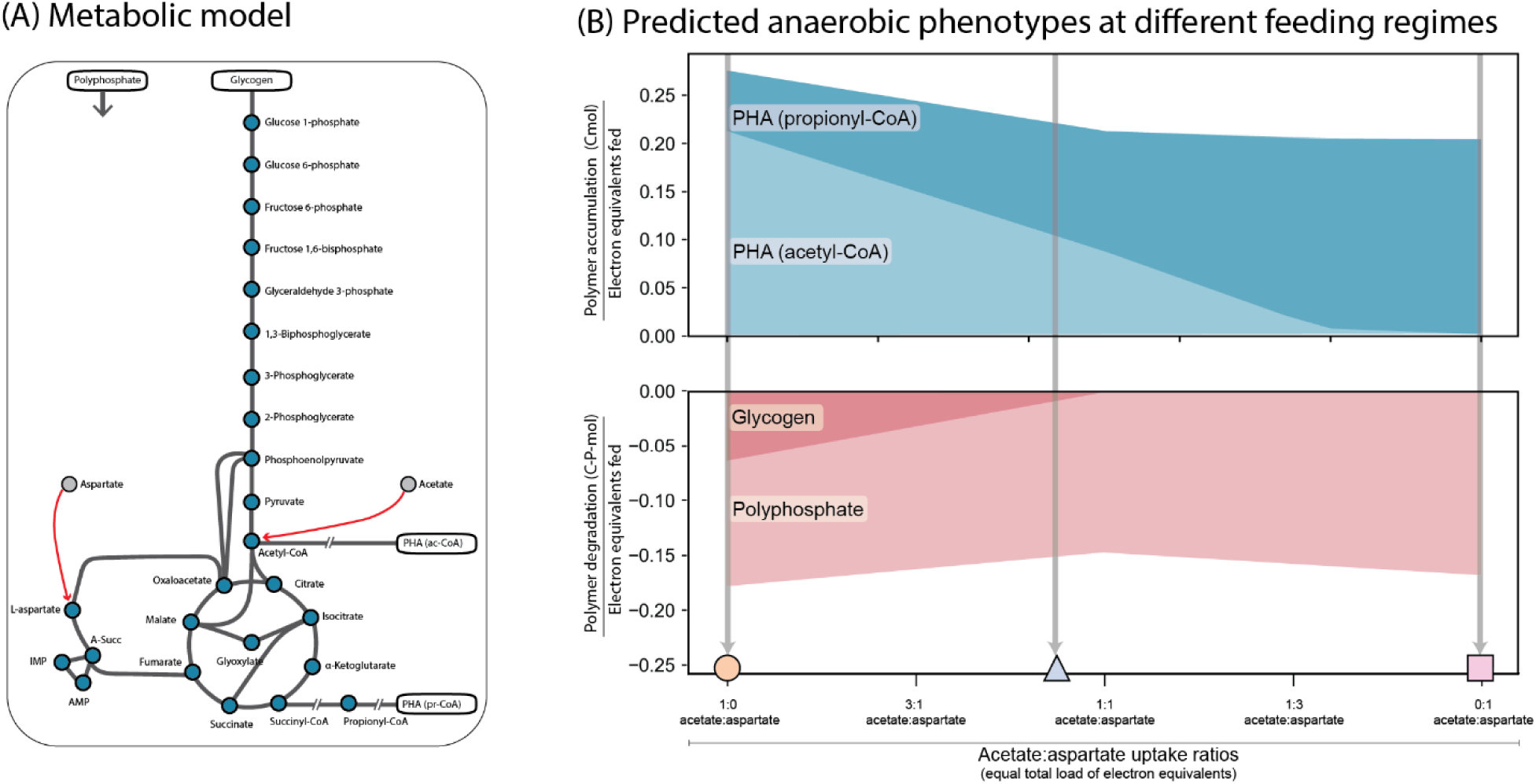
Different anaerobic stoichiometries employed for the uptake of acetate, aspartate and combined substrates. **(A)** Schematic representation of the metabolic model of “Ca. Accumulibacter” used for simulations. Metabolites are represented as filled circles connected via reactions with grey lines. Red arrows indicate the entry point for substrate uptake in central carbon metabolism. Intracellular storage polymers are represented as white boxes. **(B)** Anaerobic conversions during the uptake of acetate/aspartate at varying electron equivalent ratios simulated with cFBA. (In blue) PHA accumulation subdivided into PHAs from acetyl-CoA (C2) and from propionyl-CoA (C3). (In red) Glycogen and Polyphosphate consumption. Three specific simulations marked with symbols (circle, triangle and square) were confirmed experimentally in Figure 2.

With acetate as the sole substrate, polyP and glycogen were degraded to supply resources for PHA accumulation primarily as acetyl-CoA precursors with a smaller fraction of propionyl-CoA. As the fraction of aspartate uptake increased, less glycogen degradation was required and larger fraction of PHAs as propionyl-CoA precursors was synthesized. At an equal electron equivalence ratio of acetate to aspartate, glycogen degradation halted. In scenarios with higher aspartate fractions, glycogen degradation was not observed, and the PHA pool showed a higher dominance of propionyl-CoA precursors with a higher demand for polyphosphate degradation (Figure 1.B).

To validate the modelling results, batch tests were performed on a lab reactor enrichment culture. qFISH analysis estimated the biovolume abundance of “*Ca.* Accumulibacter” at 89 ± 3 %., while metagenomics analysis revealed the enrichment of a clade I strain closely related to “*Ca.* Accumulibacter regalis”. The enriched genome harboured the complete genetic potential required for aspartate metabolism (Figure 2.A).

**Figure 2.**
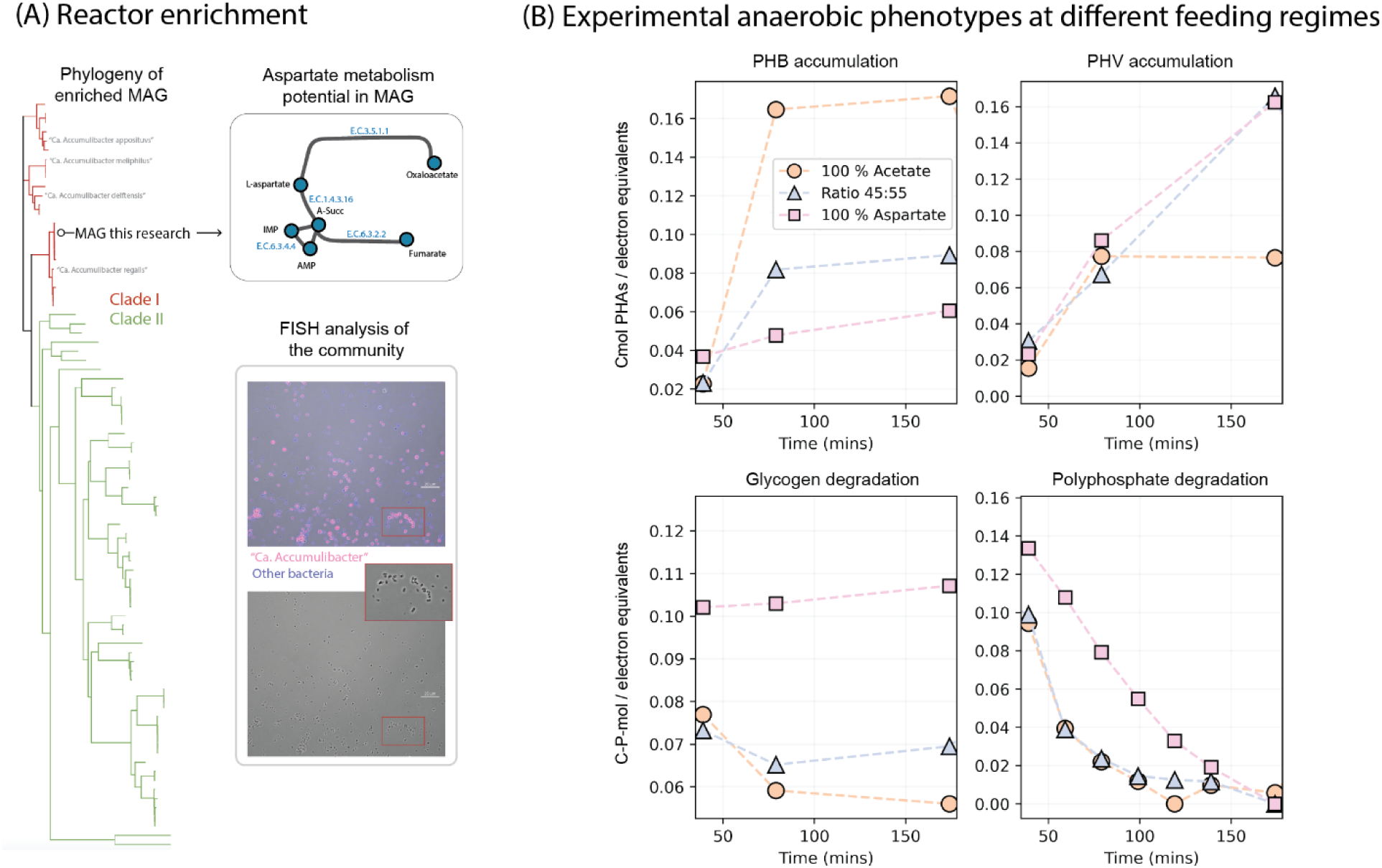
Lab culture enrichment and experimental validation of the model. **(A)** Microbial community analysis with metagenomics and FISH. Metagenomics revealed the enrichment of an MAG closely associated to Ca. Accumulibacter Clade I “Ca. Accumulibacter regalis” and harbouring all the genes necessary for aspartate metabolism. FISH image of the enrichment in which magenta colour represents the overlap of “Ca. Accumulibacter” (red) and eubacteria (blue). Bottom image of phase contrast highlighting typical morphology observed from PAOs enrichments. **(B)** Experimental validation of the anaerobic phase metabolic strategies observed during batch tests for acetate, aspartate, and mixed acetate:aspartate (45:55 electron equivalence) regimes. Distinct markers (circles, triangles, and squares) facilitate direct comparison with the modeled predictions in Figure 1.B.

Substrate compositions were evaluated in batch tests under three regimes: acetate, aspartate and a 45:55 electron equivalence ratio of acetate to aspartate consumption (regimes marked in Figure 1.B). The experimental results closely matched the predicted stoichiometries (Figure 2.B). Specifically, in the acetate-fed regime, PHAs accumulated anaerobically mainly as acetyl-CoA precursors, accompanied by the degradation of polyphosphate and glycogen. In the mixed substrate regime, PHAs accumulated as a balanced mixture of both acetyl-CoA and propionyl-CoA precursors, with lower glycogen degradation per electron equivalent consumed compared to the acetate-only regime, consistent with the predictions. Finally, in the aspartate-fed regime, PHAs accumulated with a substantial decrease in acetyl-CoA precursors. This regime required the highest polyphosphate degradation and no glycogen degradation, aligning with the predictions (See Figure 1.B and Figure 2.B for comparison).

### Metabolic and energetic balances reveal complementary strategies for the uptake of acetate and aspartate leading to enhanced growth yields

Simulations with varying ratios of acetate and aspartate in the feed revealed not only changes in internal storage polymer utilization under anaerobic conditions, but also predicted maximum growth yields per electron equivalents for each cycle (Figure 3.A). Growth on aspartate was more efficient than growth on acetate. Notably, the highest growth yield was achieved with a combination of both substrates, specifically at a 1:4 electron equivalent ratio of acetate to aspartate.

**Figure 3.**
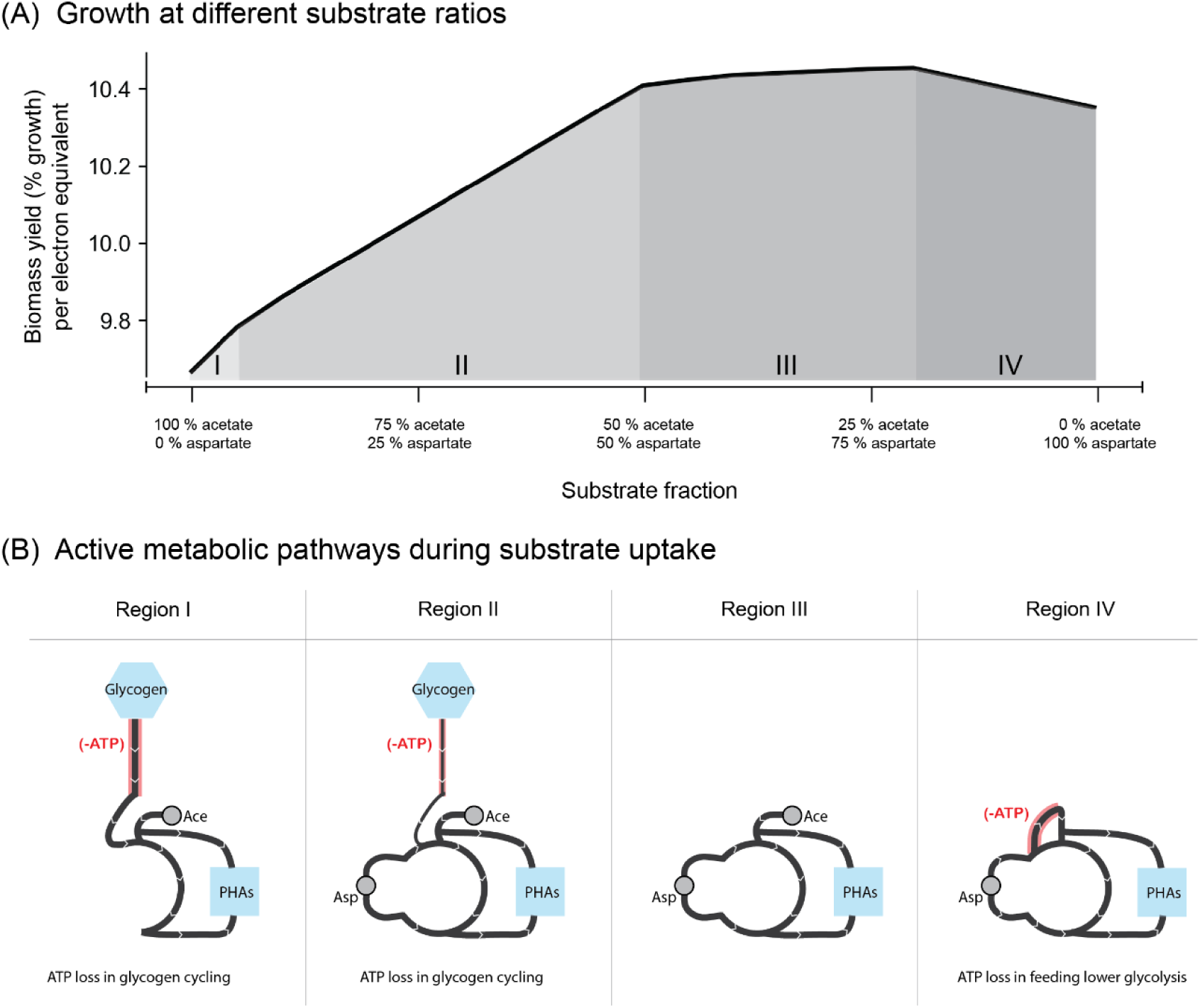
**(A)** Biomass growth yield as a duplication (% growth) in one cycle per electron equivalents fed at varying fractions of acetate aspartate. Each region represents a unique metabolic strategy employed by the model to optimize growth. **(B)** Active metabolic operation during anaerobic substrate uptake predicted via the cFBA model at each region indicated in (A). Metabolic reactions contributing to a net ATP loss of the system considering the whole EBPR cycle have been highlighted in red. Lower thickness in Region II illustrates the lower demand on glycogen degradation than in Region I.

During acetate uptake, glycogen was degraded to supply NADH necessary for PHA accumulation. The most efficient strategy for PHA accumulation reflected in the model involved glycogen degradation via glycolysis to phosphoenolpyruvate (PEP), then converting PEP to oxaloacetate (OAA) to fuel the TCA cycle. This allowed partial acetate oxidation in the right branch of the TCA cycle, producing NADH and Propionyl-CoA-type PHAs (Figure 3.B – region I). However, during the aerobic phase, this strategy required glycogen replenishment and the resulting glycolysis/gluconeogenesis operation over the cycle results in a net ATP loss, thus reducing the overall growth yield.

As the aspartate-to-acetate ratio increased, aspartate metabolism generated additional NADH via aspartate oxidase that countered the NADH requirement from glycogen degradation (Figure 3.B – region II), thus lowering the net ATP loss in the glycolysis/gluconeogenesis cycle, which improved growth yields. When the ratio reached 1:1, glycogen degradation was no longer necessary. Beyond this point, up to a 1:4 acetate-to-aspartate ratio, there were no net ATP losses, resulting in the highest growth yields (Figure 3.B – region III). In this range reciprocal benefits were observed when aspartate consumption provided NADH that benefited acetate metabolism, while acetate consumption supplied acetyl-CoA equivalents that supported aspartate metabolism (as described below).

At higher aspartate fractions, the metabolic strategy necessitated the operation of the right branch of the TCA cycle, which required acetyl-CoA equivalents. These equivalents could be supplied through acetate uptake. At insufficient acetate fractions, part of the consumed aspartate was channelled towards acetyl-CoA generation via PEP carboxykinase (PEPCK), raising the demand for ATP (Figure 3.B – region IV). This was met with an increased polyphosphate degradation, necessitating increased ATP requirements in the aerobic phase to replenish the polyphosphate pools resulting in lower growth yields.

### Synergistic effects of substrate co-consumption vary by metabolic entry point and network topology

Several substrates were incorporated into the existing metabolic model of “*Ca*. Accumulibacter”, and their co-consumption with acetate at varying ratios was simulated using cFBA, as described in the previous section. The predictions indicated that multiple substrates could support PHA accumulation without relying on reducing equivalents from glycogen degradation, which is typically required during anaerobic acetate uptake (Figure 4).

**Figure 4.**
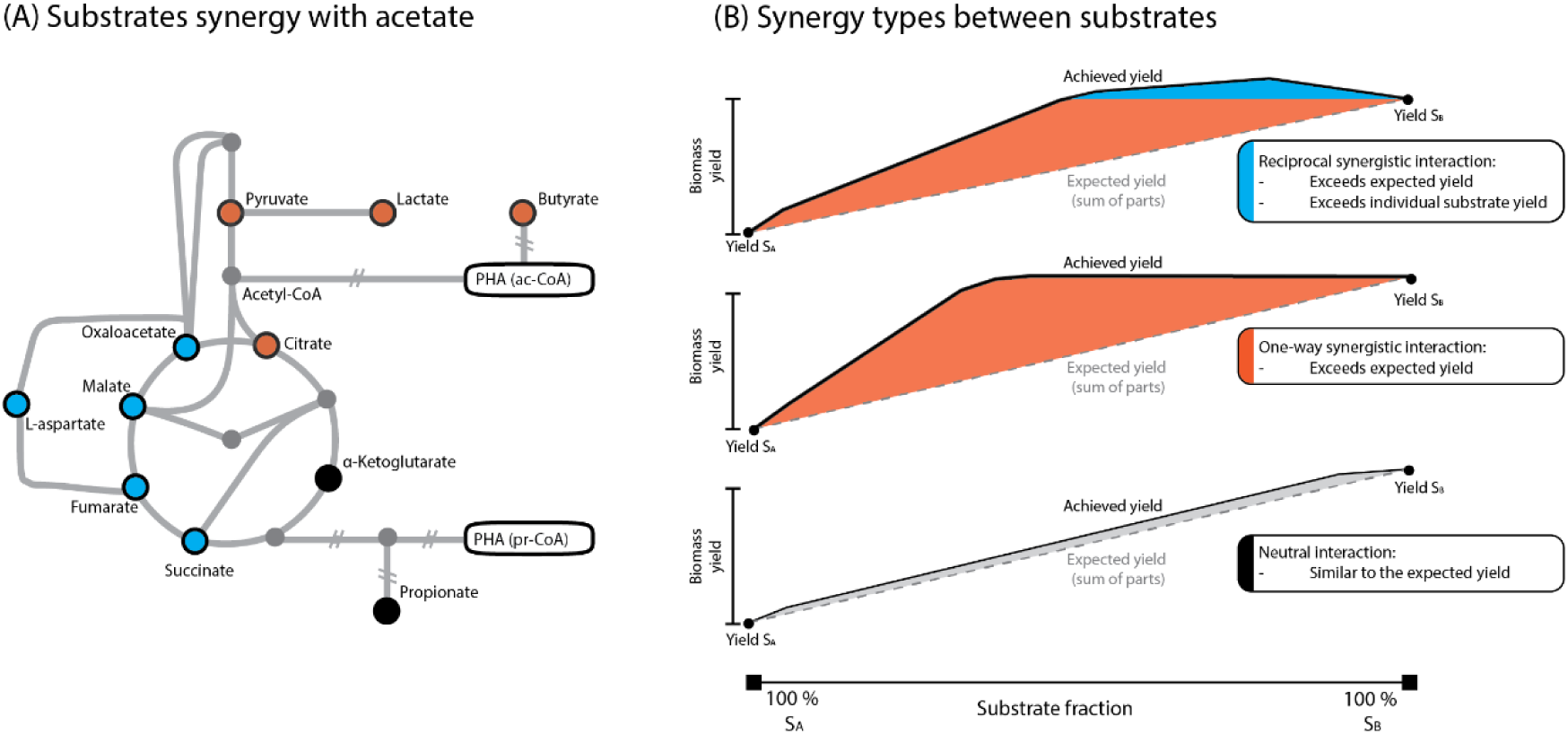
Substrates interactions when co-consumed. **(A)** Metabolic model of “Ca. Accumulibacter” highlighting the external substrates that were tested and their type of interaction when co-fed with acetate. Blue: reciprocal synergistic interaction. Orange: one-way synergistic interaction. Black: neutral interaction. **(B)** Types of interactions (substrate synergy) existing between two substrates. Top shows reciprocal synergy between acetate and aspartate in which the biomass yield can exceed that of the sum of parts and the individual maximum biomass yields. Middle panel shows a one-way synergistic interaction between acetate and lactate in which the biomass yield can exceed that of the sum of parts but not the individual maximum yield. Bottom panel shows a neutral interaction between acetate and propionate in which the biomass yield is similar to the expected from the sum of parts.

Interestingly, the co-consumption of certain substrates with acetate mirrored the reciprocal synergistic effect observed with aspartate. Specifically, these combinations led to an enhanced biomass yield per electron equivalent compared to the yield of individual substrates. We referred to these as reciprocal synergistic interactions. Substrates exhibiting this behaviour included succinate, fumarate, malate, oxaloacetate, and aspartate, all of which enter the reducing branch (left-hand side) of the TCA cycle. The metabolism of these substrates resulted in sufficient NADH production to alleviate the dependence from glycogen degradation, the main limitation during acetate uptake. Complementarily, metabolising these substrates benefited from the uptake of acetate to feed acetyl-CoA equivalents into the TCA cycle, as was the case with aspartate.

In contrast, another class of substrates co-consumed with acetate produced biomass yields greater than the sum of the individual parts but did not surpass the yield of the more favourable substrate on its own. These substrates were able to generate sufficient reducing power (NADH) to alleviate the reliance on glycogen degradation, releasing the limitation for acetate metabolism. However, they did not benefit from the additional acetate uptake, since their metabolism did not require acetyl-CoA to be fed into the TCA cycle using PEPCK or similar reactions. We classified these as one-way synergistic interactions, observed for butyrate, lactate, pyruvate and citrate.

Finally, certain substrates such as propionate and alpha-ketoglutarate resulted in biomass yields that closely matched the sum of the individual yields, with no additional gain from co-consumption. We classified these as neutral interactions, wherein the metabolic demands of these substrates closely resembled that of acetate. These substrates required similar resources (NADH and ATP) as acetate, leading to overlapping metabolic strategies that did not enhance overall growth yield.

Importantly, when thermodynamic constraints were removed from the model, allowing all reactions to operate reversibly, no significant differences were observed between the substrates (See supplementary materials). In this scenario, all co-consumed substrates resulted in similar biomass yields, producing a horizontal line in the interaction plot, demonstrating the absence of any metabolic synergy under reversible reaction conditions.

## 4. Discussion

This study demonstrates that “*Ca.* Accumulibacter” exhibits synergistic metabolic interactions when acetate is co-consumed with aspartate, leading to enhanced growth yields and reduced net ATP loss. These findings offer new insights into the metabolic mechanisms of substrate co-consumption in EBPR systems and highlight the importance of network topology in determining interaction outcomes. Below, we contextualize these results within existing research on EBPR, biological adaptation, and energy cycling while also acknowledging the limitations of the predictive models used.

Wastewater typically consists of a mix of organic substrates^17–19, 51^, and many studies have documented “Ca. Accumulibacter” as capable of consuming multiple substrates, both through genomic analysis^16, 52^ and *in situ* studies^53–55^. Our findings align with these studies, showing that co-consumption of acetate and aspartate not only is possible but also results in a metabolic interaction that optimizes cellular resource. The reciprocal synergy observed arises from a two-way release of metabolic limitations for each substrate, improving overall biomass yield (Figure 2). Specifically, acetate uptake is typically limited by anaerobic glycogen degradation, resulting in net ATP loss^9, 56^, while aspartate uptake requires ATP-consuming conversion of acetyl-CoA via PEPCK^16^. The combination of both substrates alleviates these limitations, leading to improved growth. It is noteworthy that the identified release in metabolic limitations were not dependent on potential *pmf* generation as hypothezised by Qiu et al.^16^ and thus can also explain the results obtained by Chen et al.^23^.

The energy losses associated with glycogen cycling during acetate consumption in “*Ca.* Accumulibacter” has been well documented^5, 9, 10, 57–60^, but their direct connection to physiological effects, such as growth yields, has not been previously established. Research manipulating growth rates by adjusting biomass retention time suggest that higher glycogen cycling corresponds to lower biomass yields^61–63^, though the connection to energetics and metabolism has not been discussed extensively. This behaviour can be compared to the broader context of “ATP demanding yet useful” cycles^56^. Since the ATP loss in cycling glycogen is temporally separated, this is not a futile cycle in strict sense. Yet the temporal separation of glycogen degradation and replenishment serves an adaptive purpose, allowing organisms to maintain metabolic flexibility in response to dynamic environments. This flexibility supports survival and growth under fluctuating conditions, similarly as that of apparent ATP-demanding pathways^64–67^. Further consideration of the metabolic and energetic cost/benefit of these temporally separated cycles needs to be considered.

The interaction between the metabolic operations of acetate and aspartate is complex, due to both the complexity inherent of metabolic networks^68^ and the dynamic nature of the EBPR cycle. The metabolic consequences of anaerobic uptake strategies can only be fully understood when considering the entire cycle. Most research in EBPR focuses on anaerobic processes, where the cell’s redox state is tightly constrained, limiting reaction feasibility^32^. However, without a holistic perspective on the aerobic phase, it is difficult to assess how anaerobic pathways impact overall metabolic fitness. The cFBA model employed here provides a tool to explore these interconnected processes, revealing metabolic strategies that minimize ATP losses as emergent properties of the system rather than being predefined *a priori*. Our experimental results aligned well with the model’s predictions, indicating that cFBA model successfully captured the key features of substrate interactions and energy flows. This agreement between predictions and experimental data reinforces the utility of cFBA in understanding complex metabolic behaviors.

While our model successfully identified synergistic interactions, it was not able to detect any substrate combinations that would lower biomass yields (in essence, negative interactions). This limitation stems from the model’s focus on global biomass yield optimization. Potential negative interactions might arise when dealing with substrates that activate stress responses or an overproduction of reductive potential and warrant further investigation.

Understanding the complete EBPR cycle is essential for uncovering the metabolic costs associated with PAO strategies. Studies examining different carbon substrates, such as butyrate and lactate, have shown shifts in glycogen use that may impact EBPR performance, but these effects have not been integrated into a broader understanding of biomass growth. Our results suggest that the energy wasteful use of glycogen in “*Ca.* Accumulibacter” when consuming acetate can be greatly released when co-consumed with many other substrates (amongst them, butyrate, lactate, pyruvate, citrate, oxaloacetate, malate, fumarate and succinate. See Figure 3). Further experiments are needed to uncover the metabolic effects of multiple organic substrates and how it can be employed to improve phosphorus removal.

## 5. Conclusions

- Co-consumption of acetate and aspartate by “Ca. Accumulibacter” results in synergistic metabolic interactions that improve biomass yield and reduce ATP losses.
- Acetate conversion to PHA benefits from NADH generated during aspartate metabolism, while aspartate uptake is supported by acetyl-CoA produced from acetate uptake.
- Glycogen cycling related to growth on acetate is energy demanding, co-consumption with other substrates (e.g., aspartate, succinate, fumarate) reduces this energy demand.
- A holistic consideration of the entire EBPR cycle is essential to fully understand the metabolic strategies and optimize the performance of PAOs.
- Synergistic interactions arising from metabolic optimization present an opportunity for co-utilization of carbon substrates that can be exploited to enhance the yield of bio-based processes.

## 6. Acknowledgements

**SR** was supported by the European Union’s Horizon Europe research and innovation program under the Marie Skłodowska-Curie grant agreement No 101068900. **MvL**, **SR** and **TPW** were supported by the SIAM Gravitation Grant 024.002.002, The Netherlands Organization for Scientific Research.

## References

1. Lu, H.; Oehmen, A.; Virdis, B.; Keller, J.; Yuan, Z., Obtaining highly enriched cultures of Candidatus Accumulibacter phosphates through alternating carbon sources. Water Research 2006, 40 (20), 3838–3848.

2. Páez-Watson, T.; Tomás-Martínez, S.; de Wit, R.; Keisham, S.; Tateno, H.; van Loosdrecht, M. C.; Lin, Y., Sweet Secrets: Exploring Novel Glycans and Glycoconjugates in the Extracellular Polymeric Substances of “Candidatus Accumulibacter”. ACS Es&t Water 2024, 4 (8), 3391–3399.

3. Kleikamp, H. B.; Grouzdev, D.; Schaasberg, P.; van Valderen, R.; van der Zwaan, R.; van de Wijgaart, R.; Lin, Y.; Abbas, B.; Pronk, M.; van Loosdrecht, M. C., Comparative metaproteomics demonstrates different views on the complex granular sludge microbiome. bioRxiv 2022, 2022.03. 07.483319.

4. Kleikamp, H. B.; Grouzdev, D.; Schaasberg, P.; van Valderen, R.; van der Zwaan, R.; van de Wijgaart, R.; Lin, Y.; Abbas, B.; Pronk, M.; van Loosdrecht, M. C., Metaproteomics, metagenomics and 16S rRNA sequencing provide different perspectives on the aerobic granular sludge microbiome. Water research 2023, 246, 120700.

5. Mino, T.; Arun, V.; Tsuzuki, Y.; Matsuo, T., Effect of phosphorus accumulation on acetate metabolism in the biological phosphorus removal process. In Biological phosphate removal from wastewaters, Elsevier: 1987; pp 27–38.

6. Kortstee, G. J.; Appeldoorn, K. J.; Bonting, C. F.; van Niel, E. W.; van Veen, H. W., Biology of polyphosphate-accumulating bacteria involved in enhanced biological phosphorus removal. FEMS microbiology reviews 1994, 15 (2-3), 137–153.

7. Smolders, G.; Van der Meij, J.; Van Loosdrecht, M.; Heijnen, J., Model of the anaerobic metabolism of the biological phosphorus removal process: stoichiometry and pH influence. Biotechnology and bioengineering 1994, 43 (6), 461–470.

8. Smolders, G.; Van der Meij, J.; Van Loosdrecht, M.; Heijnen, J., Stoichiometric model of the aerobic metabolism of the biological phosphorus removal process. Biotechnology and bioengineering 1994, 44 (7), 837–848.

9. Smolders, G.; Van der Meij, J.; Van Loosdrecht, M.; Heijnen, J., A structured metabolic model for anaerobic and aerobic stoichiometry and kinetics of the biological phosphorus removal process. Biotechnology and Bioengineering 1995, 47 (3), 277–287.

10. Van Loosdrecht, M.; Smolders, G.; Kuba, T.; Heijnen, J., Metabolism of micro-organisms responsible for enhanced biological phosphorus removal from wastewater, Use of dynamic enrichment cultures. Antonie van Leeuwenhoek 1997, 71 (1-2), 109–116.

11. Mino, T.; Van Loosdrecht, M.; Heijnen, J., Microbiology and biochemistry of the enhanced biological phosphate removal process. Water Research 1998, 32 (11), 3193–3207.

12. Welles, L.; Tian, W.; Saad, S.; Abbas, B.; Lopez-Vazquez, C.; Hooijmans, C.; Van Loosdrecht, M.; Brdjanovic, D., Accumulibacter clades Type I and II performing kinetically different glycogen-accumulating organisms metabolisms for anaerobic substrate uptake. Water research 2015, 83, 354–366.

13. Pijuan, M.; Saunders, A. M.; Guisasola, A.; Baeza, J. A.; Casas, C.; Blackall, L., Enhanced biological phosphorus removal in a sequencing batch reactor using propionate as the sole carbon source. Biotechnology and bioengineering 2004, 85 (1), 56–67.

14. Elahinik, A.; Li, L.; Pabst, M.; Abbas, B.; Xevgenos, D.; van Loosdrecht, M. C.; Pronk, M., Aerobic granular sludge phosphate removal using glucose. Water Research 2023, 247, 120776.

15. Ziliani, A.; Bovio-Winkler, P.; Cabezas, A.; Etchebehere, C.; Garcia, H. A.; López-Vázquez, C. M.; Brdjanovic, D.; van Loosdrecht, M. C.; Rubio-Rincón, F. J., Putative metabolism of Ca. Accumulibacter via the utilization of glucose. Water Research 2023, 229, 119446.

16. Qiu, G.; Liu, X.; Saw, N. M. M. T.; Law, Y.; Zuniga-Montanez, R.; Thi, S. S.; Ngoc Nguyen, T. Q.; Nielsen, P. H.; Williams, R. B.; Wuertz, S., Metabolic traits of Candidatus Accumulibacter clade IIF strain SCELSE-1 using amino acids as carbon sources for enhanced biological phosphorus removal. Environmental Science & Technology 2019, 54 (4), 2448–2458.

17. Huang, M.-h.; Li, Y.-m.; Gu, G.-w., Chemical composition of organic matters in domestic wastewater. Desalination 2010, 262 (1-3), 36–42.

18. Bengtsson, S.; Hallquist, J.; Werker, A.; Welander, T., Acidogenic fermentation of industrial wastewaters: Effects of chemostat retention time and pH on volatile fatty acids production. Biochemical Engineering Journal 2008, 40 (3), 492–499.

19. Qiu, G.; Zuniga-Montanez, R.; Law, Y.; Thi, S. S.; Nguyen, T. Q. N.; Eganathan, K.; Liu, X.; Nielsen, P. H.; Williams, R. B.; Wuertz, S., Polyphosphate-accumulating organisms in full-scale tropical wastewater treatment plants use diverse carbon sources. Water research 2019, 149, 496–510.

20. Gebremariam, S. Y.; Beutel, M. W.; Christian, D.; Hess, T. F., Effects of glucose on the performance of enhanced biological phosphorus removal activated sludge enriched with acetate. Bioresource Technology 2012, 121, 19–24.

21. Rubio-Rincón, F. J.; Welles, L.; Lopez-Vazquez, C. M.; Abbas, B.; van Loosdrecht, M. C.; Brdjanovic, D., Effect of lactate on the microbial community and process performance of an EBPR system. Frontiers in microbiology 2019, 10, 125.

22. Yuan, Q.; Sparling, R.; Lagasse, P.; Lee, Y.; Taniguchi, D.; Oleszkiewicz, J., Enhancing biological phosphorus removal with glycerol. Water Science and Technology 2010, 61 (7), 1837–1843.

23. Chen, L.; Wei, G.; Zhang, Y.; Wang, K.; Wang, C.; Deng, X.; Li, Y.; Xie, X.; Chen, J.; Huang, F., Candidatus Accumulibacter use fermentation products for enhanced biological phosphorus removal. Water Research 2023, 246, 120713.

24. Nielsen, J., Systems biology of metabolism. Annual review of biochemistry 2017, 86 (1), 245–275.

25. Orth, J. D.; Thiele, I.; Palsson, B. Ø., What is flux balance analysis? Nature biotechnology 2010, 28 (3), 245–248.

26. Mahadevan, R.; Edwards, J. S.; Doyle III, F. J., Dynamic flux balance analysis of diauxic growth in Escherichia coli. Biophysical journal 2002, 83 (3), 1331–1340.

27. Sarkar, D.; Mueller, T. J.; Liu, D.; Pakrasi, H. B.; Maranas, C. D., A diurnal flux balance model of Synechocystis sp. PCC 6803 metabolism. PLoS computational biology 2019, 15 (1), e1006692.

28. Liu, L.; Bockmayr, A., Regulatory dynamic enzyme-cost flux balance analysis: A unifying framework for constraint-based modeling. Journal of Theoretical Biology 2020, 501, 110317.

29. Rügen, M.; Bockmayr, A.; Steuer, R., Elucidating temporal resource allocation and diurnal dynamics in phototrophic metabolism using conditional FBA. Scientific reports 2015, 5, 15247.

30. Páez-Watson, T.; van Loosdrecht, M. C.; Wahl, S. A., Predicting the impact of temperature on metabolic fluxes using resource allocation modelling: Application to polyphosphate accumulating organisms. Water Research 2023, 228, 119365.

31. Páez-Watson, T.; Hernández Medina, R.; Vellekoop, L.; van Loosdrecht, M. C.; Wahl, S. A., Conditional Flux Balance Analysis (cFBA) Toolbox for python: application to research metabolism in cyclic environments. Bioinformatics Advances 2024, vbae174.

32. Páez-Watson, T.; van Loosdrecht, M. C.; Wahl, S. A., From metagenomes to metabolism: Systematically assessing the metabolic flux feasibilities for “Candidatus Accumulibacter” species during anaerobic substrate uptake. Water Research 2024, 250, 121028.

33. Gommers, P.; Van Schie, B.; Van Dijken, J.; Kuenen, J., Biochemical limits to microbial growth yields: an analysis of mixed substrate utilization. Biotechnology and bioengineering 1988, 32 (1), 86–94.

34. Oehmen, A.; Zeng, R. J.; Keller, J.; Yuan, Z., Modeling the Aerobic Metabolism of Polyphosphate-Accumulating Organisms Enriched with Propionate as a Carbon Source. Water Environment Research 2007, 79 (13), 2477–2486.

35. Winkler, M.-K.; Bassin, J.; Kleerebezem, R.; De Bruin, L.; Van den Brand, T.; Van Loosdrecht, M., Selective sludge removal in a segregated aerobic granular biomass system as a strategy to control PAO– GAO competition at high temperatures. Water research 2011, 45 (11), 3291–3299.

36. Amann, R. I.; Binder, B. J.; Olson, R. J.; Chisholm, S. W.; Devereux, R.; Stahl, D., Combination of 16S rRNA-targeted oligonucleotide probes with flow cytometry for analyzing mixed microbial populations. Applied and environmental microbiology 1990, 56 (6), 1919–1925.

37. Daims, H.; Brühl, A.; Amann, R.; Schleifer, K.-H.; Wagner, M., The domain-specific probe EUB338 is insufficient for the detection of all Bacteria: development and evaluation of a more comprehensive probe set. Systematic and applied microbiology 1999, 22 (3), 434–444.

38. Petriglieri, F.; Singleton, C. M.; Kondrotaite, Z.; Dueholm, M. K.; McDaniel, E. A.; McMahon, K. D.; Nielsen, P. H., Reevaluation of the Phylogenetic Diversity and Global Distribution of the Genus “Candidatus Accumulibacter”. mSystems 2022, 7 (3), e00016–22.

39. Daims, H.; Lücker, S.; Wagner, M., Daime, a novel image analysis program for microbial ecology and biofilm research. Environmental microbiology 2006, 8 (2), 200–213.

40. Andrews, S., FastQC: a quality control tool for high throughput sequence data. Cambridge, United Kingdom: 2010.

41. Chen, S., Ultrafast one-pass FASTQ data preprocessing, quality control, and deduplication using fastp. Imeta 2023, 2 (2), e107.

42. Wood, D. E.; Lu, J.; Langmead, B., Improved metagenomic analysis with Kraken 2. Genome biology 2019, 20, 1–13.

43. Nurk, S.; Meleshko, D.; Korobeynikov, A.; Pevzner, P. A., metaSPAdes: a new versatile metagenomic assembler. Genome research 2017, 27 (5), 824–834.

44. Kang, D. D.; Li, F.; Kirton, E.; Thomas, A.; Egan, R.; An, H.; Wang, Z., MetaBAT 2: an adaptive binning algorithm for robust and efficient genome reconstruction from metagenome assemblies. PeerJ 2019, 7, e7359.

45. Parks, D. H.; Imelfort, M.; Skennerton, C. T.; Hugenholtz, P.; Tyson, G. W., CheckM: assessing the quality of microbial genomes recovered from isolates, single cells, and metagenomes. Genome research 2015, 25 (7), 1043–1055.

46. Chaumeil, P.-A.; Mussig, A. J.; Hugenholtz, P.; Parks, D. H., GTDB-Tk v2: memory friendly classification with the genome taxonomy database. Bioinformatics 2022, 38 (23), 5315–5316.

47. Johnson, L. S.; Eddy, S. R.; Portugaly, E., Hidden Markov model speed heuristic and iterative HMM search procedure. BMC bioinformatics 2010, 11, 1–8.

48. Edgar, R. C., MUSCLE: a multiple sequence alignment method with reduced time and space complexity. BMC bioinformatics 2004, 5, 1–19.

49. Kozlov, A. M.; Darriba, D.; Flouri, T.; Morel, B.; Stamatakis, A., RAxML-NG: a fast, scalable and user-friendly tool for maximum likelihood phylogenetic inference. Bioinformatics 2019, 35 (21), 4453–4455.

50. Letunic, I.; Bork, P., Interactive Tree of Life (iTOL) v6: recent updates to the phylogenetic tree display and annotation tool. Nucleic Acids Research 2024, gkae268.

51. Toja Ortega, S.; Pronk, M.; de Kreuk, M. K., Anaerobic hydrolysis of complex substrates in full-scale aerobic granular sludge: enzymatic activity determined in different sludge fractions. Applied Microbiology and Biotechnology 2021, 105 (14), 6073–6086.

52. Oyserman, B. O.; Noguera, D. R.; del Rio, T. G.; Tringe, S. G.; McMahon, K. D., Metatranscriptomic insights on gene expression and regulatory controls in Candidatus Accumulibacter phosphatis. The ISME journal 2016, 10 (4), 810–822.

53. Kong, Y.; Nielsen, J. L.; Nielsen, P. H., Microautoradiographic study of Rhodocyclus-related polyphosphate-accumulating bacteria in full-scale enhanced biological phosphorus removal plants. Applied and environmental microbiology 2004, 70 (9), 5383–5390.

54. Nguyen, H. T. T.; Kristiansen, R.; Vestergaard, M.; Wimmer, R.; Nielsen, P. H., Intracellular accumulation of glycine in polyphosphate-accumulating organisms in activated sludge, a novel storage mechanism under dynamic anaerobic-aerobic conditions. Applied and Environmental Microbiology 2015, 81 (14), 4809–4818.

55. Roy, S.; Guanglei, Q.; Zuniga-Montanez, R.; Williams, R. B.; Wuertz, S., Recent advances in understanding the ecophysiology of enhanced biological phosphorus removal. Current Opinion in Biotechnology 2021, 67, 166–174.

56. Sharma, A. K.; Khandelwal, R.; Wolfrum, C., Futile cycles: Emerging utility from apparent futility. Cell Metabolism 2024.

57. da Silva, L. G.; Gamez, K. O.; Gomes, J. C.; Akkermans, K.; Welles, L.; Abbas, B.; van Loosdrecht, M. C.; Wahl, S. A., Revealing the Metabolic Flexibility of “Candidatus Accumulibacter phosphatis” through Redox Cofactor Analysis and Metabolic Network Modeling. Applied Environmental Microbiology 2020, 86 (24).

58. Brdjanovic, D.; Van Loosdrecht, M.; Hooijmans, C.; Mino, T.; Alaerts, G.; Heijnen, J., Effect of polyphosphate limitation on the anaerobic metabolism of phosphorus-accumulating microorganisms. Applied microbiology biotechnology 1998, 50 (2), 273–276.

59. Brdjanovic, D.; van Loosdrecht, M. C.; Hooijmans, C. M.; Mino, T.; Alaerts, G. J.; Heijnen, J. J., Bioassay for glycogen determination in biological phosphorus removal systems. Water Science and Technology 1998, 37 (4-5), 541–547.

60. Oehmen, A.; Carvalho, G.; Lopez-Vazquez, C.; Van Loosdrecht, M.; Reis, M., Incorporating microbial ecology into the metabolic modelling of polyphosphate accumulating organisms and glycogen accumulating organisms. Water research 2010, 44 (17), 4992–5004.

61. Rodrigo, M.; Seco, A.; Ferrer, J.; Penya-Roja, J., The effect of sludge age on the deterioration of the enhanced biological phosphorus removal process. Environmental technology 1999, 20 (10), 1055–1063.

62. Whang, L. M.; Park, J. K., Competition between polyphosphate-and glycogen-accumulating organisms in enhanced-biological-phosphorus-removal systems: Effect of temperature and sludge age. Water Environment Research 2006, 78 (1), 4–11.

63. Onnis-Hayden, A.; Majed, N.; Li, Y.; Rahman, S. M.; Drury, D.; Risso, L.; Gu, A. Z., Impact of solid residence time (SRT) on functionally relevant microbial populations and performance in full-scale enhanced biological phosphorus removal (EBPR) systems. Water Environment Research 2020, 92 (3), 389–402.

64. Chitraju, C.; Mejhert, N.; Haas, J. T.; Diaz-Ramirez, L. G.; Grueter, C. A.; Imbriglio, J. E.; Pinto, S.; Koliwad, S. K.; Walther, T. C.; Farese, R. V., Triglyceride synthesis by DGAT1 protects adipocytes from lipid-induced ER stress during lipolysis. Cell metabolism 2017, 26 (2), 407–418. e3.

65. Sharma, A. K.; Wolfrum, C., Lipid cycling isn’t all futile. Nature metabolism 2023, 5 (4), 540–541.

66. Zhang, Y.; Li, C.; Li, H.; Song, Y.; Zhao, Y.; Zhai, L.; Wang, H.; Zhong, R.; Tang, H.; Zhu, D., miR-378 activates the pyruvate-PEP futile cycle and enhances lipolysis to ameliorate obesity in mice. EBioMedicine 2016, 5, 93–104.

67. Adolfsen, K. J.; Brynildsen, M. P., Futile cycling increases sensitivity toward oxidative stress in Escherichia coli. Metabolic engineering 2015, 29, 26–35.

68. Liu, N.; Santala, S.; Stephanopoulos, G., Mixed carbon substrates: a necessary nuisance or a missed opportunity? Current opinion in biotechnology 2020, 62, 15–21.

