## Supplementary Materials for "Metabolic implications for dual substrate growth in “*Candidatus* Accumulibacter”"

### S.M.1. Conditional Flux Balance Analysis (cFBA) model architecture

For access to the files and complete information of the simulations performed, we refer the reader to our GitHub: https://github.com/TP-Watson/PAOs_co-substrates_cFBA. Here we detail variables used in the model and the basic structure and algorithm applied in our research.

The starting point for the metabolic model is the Stoichiometric Matrix (**S**). This represents the relationship between metabolites interconnected by reactions of the network. This matrix is represented in *Table S.1.* All SBML models generated together with the excel files containing the basic structure for each simulation are available at: https://github.com/TP-Watson/PAOs_co-substrates_cFBA.

For each simulation in cFBA, we calculated the flux of feed of acetate, aspartate or a combination in order to maintain always the same chemical oxygen demand (COD), also known as degree of reduction or electron equivalence. For that, the quota compound definitions were used, in which we can determine the exact level of a given *imbalanced metabolite* at a specific moment in time. As an example, below we show the quotas and results from simulations used as reference on acetate only and on aspartate only.

```
# Quotas for the model
quotas = [
#    ['type',      'metabolite', time, value]
    ['equality', 'Acetate',    0,    0.04],
    ['equality', 'Acetate',    1,    0],
    ['equality', 'Asp_ext',    0,    0],
    ['equality', 'Asp_ext',    1,    0],
    ['min',      'Glyc',       0,    0.012],
    ['min',      'PolyP',      0,    0.1],
    ['min',      'PHB',        0,    0.006],
    ['min',      'PH2MV',      0,    0.006],
]
```

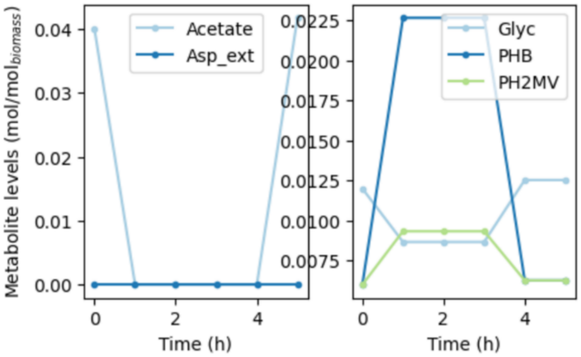

***Figure S1***. *Quotas and simulation results when performing cFBA on only acetate.*

```
quotas = [
#    ['type',      'metabolite', time, value]
    ['equality', 'Acetate',    0,    0],
    ['equality', 'Acetate',    1,    0],
    ['equality', 'Asp_ext',    0,    0.04],
    ['equality', 'Asp_ext',    1,    0],
    ['min',      'Glyc',       0,    0.012],
    ['min',      'PolyP',      0,    0.1],
    ['min',      'PHB',        0,    0.006],
    ['min',      'PH2MV',      0,    0.006],
]
```

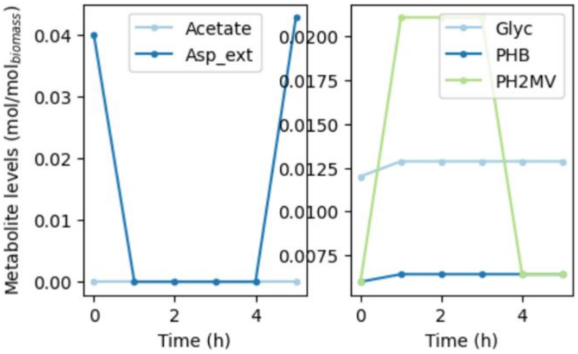

***Figure S2***. *Quotas and simulation results when performing cFBA on only acetate.*

**Table S1.** Metabolic model stoichiometry and reaction bounds applied in cFBA.

| Reaction name | Equation | Lower bound | **Upper bound** | Time dependence |
|---|---|---|---|---|
| Ac_1 | 1 Acetate + 1 ATP ---> 1 AcCoA | 0 | 1000 | - |
| Glyc_S | 1 G6P + 1 ATP ---> 1 Glyc | 0 | 1000 | - |
| Glyc_D | 1 Glyc ---> 1 G6P | 0 | 1000 | - |
| EMP1 | 1 G6P ---> 1 F6P | -1000 | 1000 | - |
| EMP2 | 1 F6P + 1 ATP ---> 1 F16P | 0 | 1000 | - |
| EMP3 | 1 F16P ---> 1 DHAP + 1 GA3P | -1000 | 1000 | - |
| EMP4 | 1 DHAP ---> 1 GA3P | -1000 | 1000 | - |
| EMP5 | 1 GA3P ---> 1 PG3 + 1 NADH | -1000 | 1000 | - |
| EMP6 | 1 PG3 ---> 1 PEP + 1 ATP | -1000 | 1000 | - |
| EMP7 | 1 PEP ---> 1 Pyr + 1 ATP | 0 | 1000 | - |
| FBPase | 1 F16P ---> 1 F6P | 0 | 1000 | - |
| TCA1 | 1 Pyr ---> 1 AcCoA + 1 NADH + 1 CO2 | 0 | 1000 | - |
| TCA2 | 1 AcCoA + 1 OAA ---> 1 Cit | -1000 | 1000 | - |
| TCA3 | 1 Cit ---> 1 iCit | -1000 | 1000 | - |
| TCA4 | 1 iCit ---> 1 aKG + 1 NADH + 1 CO2 | -1000 | 1000 | - |
| TCA5 | 1 aKG ---> 1 SuccCoA + 1 NADH + 1 CO2 | -1000 | 1000 | - |
| TCA6 | 1 SuccCoA ---> 1 Succ + 1 ATP | -1000 | 1000 | - |
| TCA7 | 1 Succ ---> 1 Fum + 1 FADH2 | -1000 | 1000 | - |

| | | | | |
|---|---|---|---|---|
| TCA8 | 1 Fum ---> 1 Mal | -1000 | 1000 | - |
| TCA9 | 1 OAA + 1 NADH ---> 1 Mal | *variable* | 1000 | Reversible only aerobically |
| NADH_FADH | 1 NADH ---> 1 FADH2 | 0 | 1000 | - |
| Succ_prop | 1 SuccCoA ---> 1 PrCoA + 1 CO2 | -1000 | 1000 | - |
| GS1 | 1 iCit ---> 1 Succ + 1 Glx | 0 | 1000 | - |
| GS2 | 1 AcCoA + 1 Glx ---> 1 Mal | 0 | 1000 | - |
| PEPCK | 1 OAA + 1 ATP ---> 1 PEP + 1 CO2 | 0 | 1000 | - |
| PEPC | 1 PEP + 1 CO2 ---> 1 OAA | 0 | 1000 | - |
| ETC_NADH | 1 NADH ---> 1.85 ATP | 0 | *variable* | Active only aerobically |
| ETC_FADH2 | 1 FADH2 ---> 1.35 ATP | 0 | *variable* | Active only aerobically |
| PolyP_S | 1 ATP ---> 1 PolyP | 0 | 1000 | - |
| PolyP_D | 1 PolyP ---> 1 ATP | 0 | 1000 | - |
| PHB_S | 2 AcCoA + 1 NADH ---> 1 PHB | 0 | 1000 | - |
| PHB_D | 1 PHB + 2 ATP ---> 2 AcCoA + 1 NADH | 0 | 1000 | - |
| PH2MV_S | 2 PrCoA + 1 NADH ---> 1 PH2MV | 0 | 1000 | - |
| PH2MV_D | 1 PH2MV + 2 ATP ---> 2 PrCoA + 1 NADH | 0 | 1000 | - |
| BM | 0.635 AcCoA + 1.7 ATP ---> 0.89 NADH + 0.27 CO2 + 1 Biomass | 0 | *variable* | Active only aerobically |
| CO2_im | ---> 1 CO2 | -1000 | 1000 | - |
| Vcomp | 1 Acetate ---> | 0 | *variable* | Active during feed |
| Ac_Feed | ---> 1 Acetate | 0 | *variable* | Active during feed |

| | | | | |
|---|---|---|---|---|
| S_Feed | ---> 1 S_ext | 0 | ***variable*** | Active during feed |
| S_up | 1 ATP + 1 S_ext ---> 1 aKG | 0 | ***variable*** | Active during feed |

### S.M.2. Co-substrates tests with no thermodynamic constraints

For access to the files and complete information of the simulations performed, we refer the reader to our GitHub: https://github.com/TP-Watson/PAOs_co-substrates_cFBA. The simulations performed for this analysis are specifically in the folder named '*cFBA co substrates tests*'.

When testing multiple substrates uptake, we performed cFBA (optimizing growth in a cycle) with varying amounts of acetate in combination with another substrate. As mentioned in the previous section, we always calculated the same electron equivalence to be the baseline for the substrate composition. We employed a mix of acetate with: alpha-keto glutarate, aspartate, butyrate, citrate, fumarate, glutamate, lactate, malate, oxaloacetate, propionate, pyruvate and succinate. For each substrate combination, we performed simulations ranging from acetate-only towards alternative substrate-only. The resulting growth yield obtained in each condition was recorded and its change against the substrate fractions revealed the types of interaction existing in the model.

#### Original model with thermodynamic constraints

The original model with the constraints described in *Table S1* led to different types of substrate interactions such as: synergistic interaction, one-sided synergy and neutral interactions. These are represented in the main text in Figure 4. The resulting growth yields for each substrate combined with acetate is represented in Figure S3.

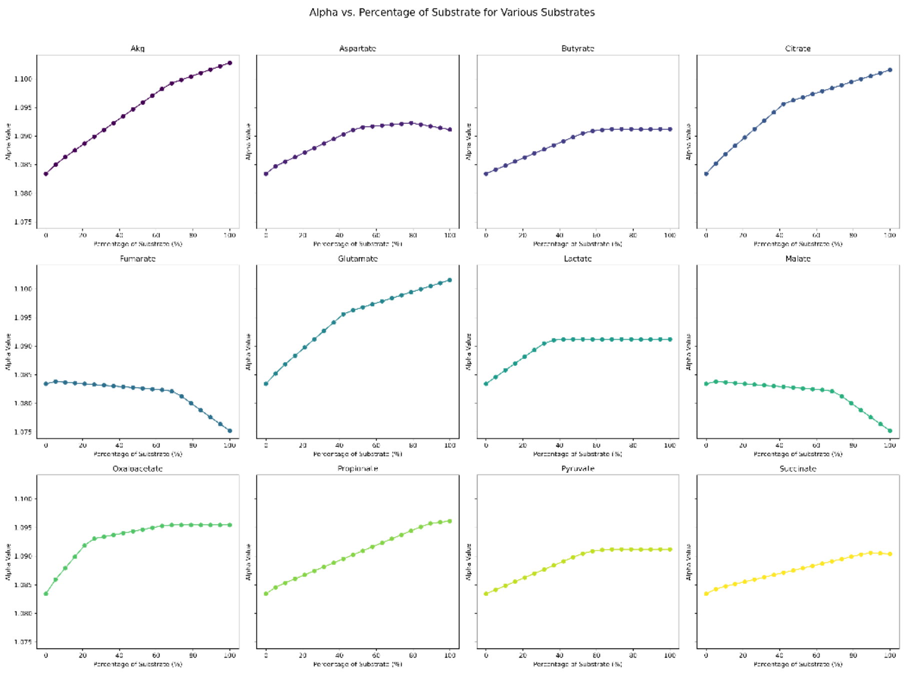

***Figure S3**. **Original model:** Resulting growth yields in each simulation with cFBA. Each dot represents the yield from a simulation in which a combination of acetate with a fraction of alternative substrate were fed.*

#### Original model allowing PEPC – PEPCK cycle

The same simulations were performed on the model, but in this time we released the thermodynamic limitation that makes the reactions 'PEPC' and 'PEPCK' irreversible (*i.e.* now tested both with lower and upper bounds of 1000). This allows a potential thermodynamic impossible cycle that can generate ATP. Nevertheless, it was tested to observe the effect on the response variable: growth yields. The results are represented in Figure S4. The release of this constraint broke the synergistic effect observed when acetate was co-fed with alpha-keto glutarate, aspartate, butyrate, citrate, glutamate, lactate, oxaloacetate, propionate, pyruvate and succinate. Only fumarate and malate maintained some type of interactions, indicating another potential mechanism at play. Importantly, the effect of releasing the PEPCK-PEPC constraint resulted in the highest growth yields on acetate, proving that acetate uptake is limited by the ATP demanding reaction required to feed the TCA cycle to generate reducing equivalents (as explained in the main text).

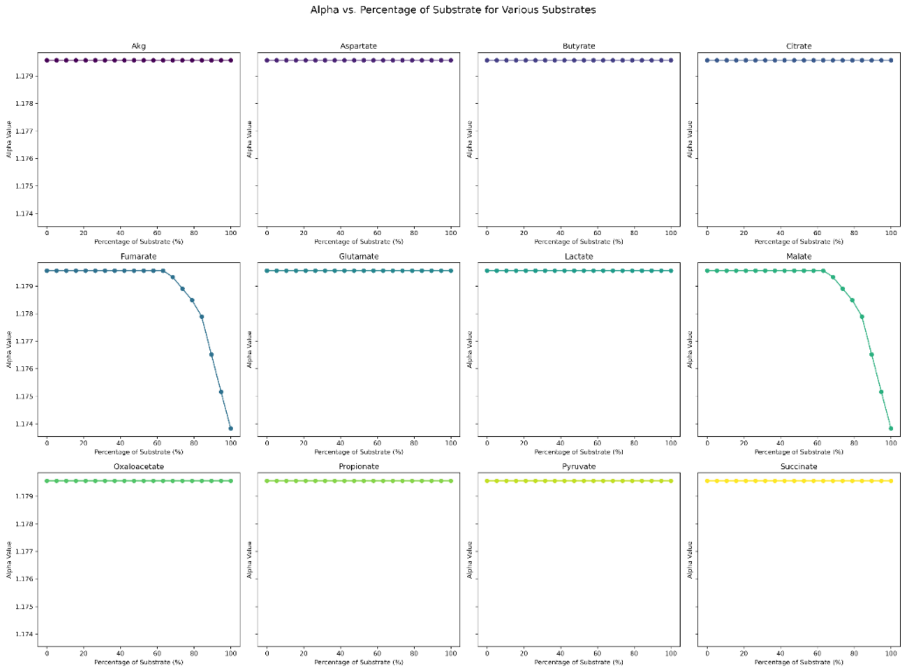

***Figure S4*. *PEPCK and PEPC reversible:*** *Resulting growth yields in each simulation with cFBA. Each dot represents the yield from a simulation in which a combination of acetate with a fraction of alternative substrate were fed. This model ignores the thermodynamic constraints of PEPCK and PEPC, allowing them both to be reversible.*

### S.M.3. FISH images from reactor enrichment

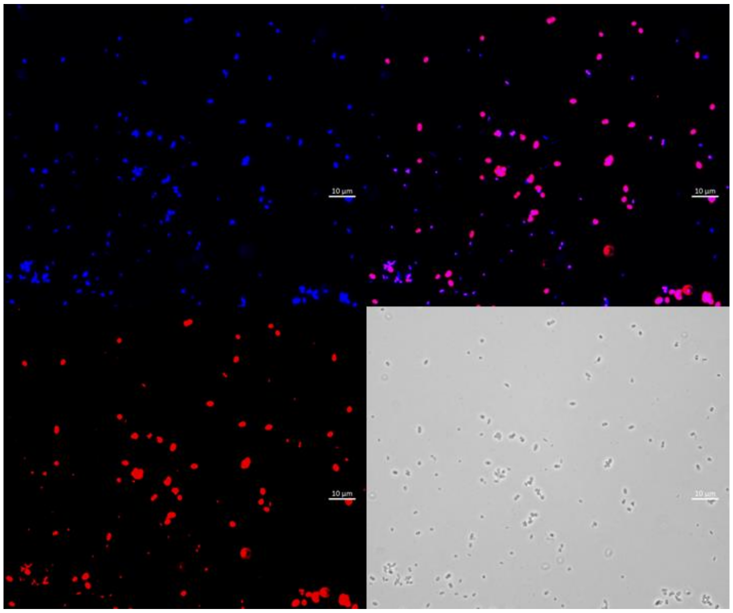

***Figure S5**. **FISH staining:** Microscopic FISH image of samples from the reactor. Blue panel represents eubacteria using a blend of EUB338, EUB338-II, and EUB338-III probes. Red panel represents "Ca. Accumulibacter" employing mixtures of the probes Acc1011, Acc471, Acc471_2, Acc635, Acc470. Magenta represents the overlap of both eubacteria and "Ca.* Accumulibacter" cells. Phase contrast shows the morphology of the cells stained.

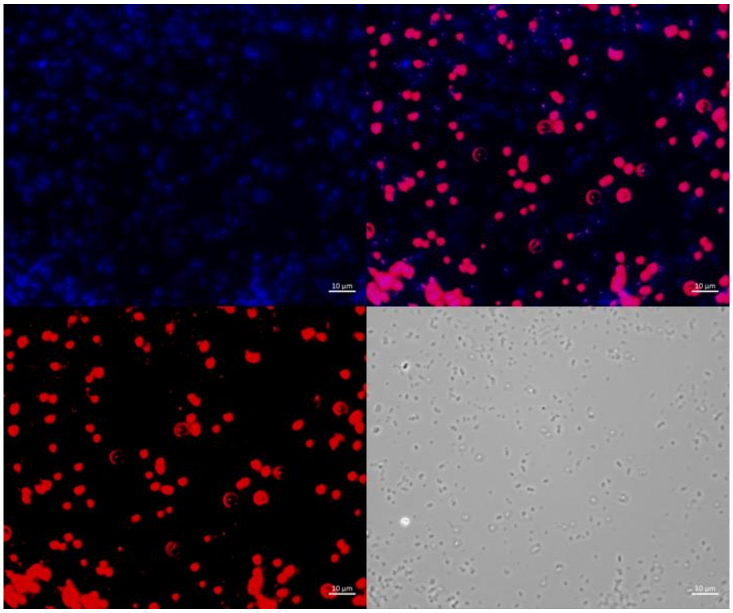

***Figure S6**. **FISH staining:** Microscopic FISH image of samples from the reactor. Blue panel represents eubacteria using a blend of EUB338, EUB338-II, and EUB338-III probes. Red panel represents "Ca. Accumulibacter" employing mixtures of the probes Acc1011, Acc471, Acc471_2, Acc635, Acc470. Magenta represents the overlap of both eubacteria and "Ca.* Accumulibacter" cells. Phase contrast shows the morphology of the cells stained.

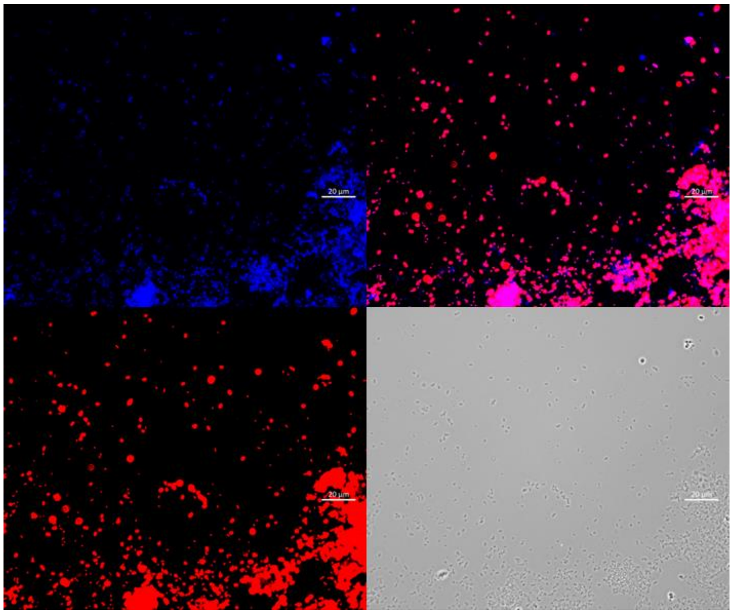

***Figure S7*. *FISH staining:*** *Microscopic FISH image of samples from the reactor. Blue panel represents eubacteria using a blend of EUB338, EUB338-II, and EUB338-III probes. Red panel represents "Ca. Accumulibacter" employing mixtures of the probes Acc1011, Acc471, Acc471_2, Acc635, Acc470. Magenta represents the overlap of both eubacteria and "Ca.* Accumulibacter" cells. Phase contrast shows the morphology of the cells stained.

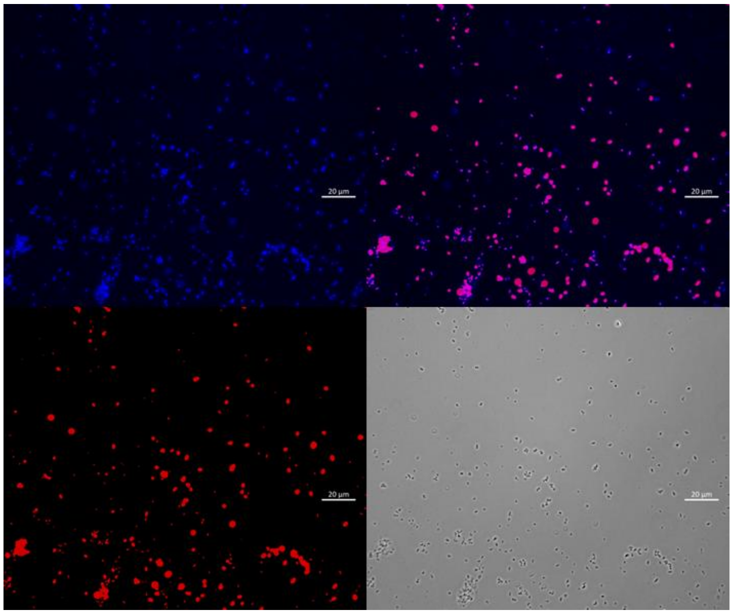

***Figure S8*. *FISH staining:*** *Microscopic FISH image of samples from the reactor. Blue panel represents eubacteria using a blend of EUB338, EUB338-II, and EUB338-III probes. Red panel represents "Ca. Accumulibacter" employing mixtures of the probes Acc1011, Acc471, Acc471_2, Acc635, Acc470. Magenta represents the overlap of both eubacteria and "Ca.* Accumulibacter" cells. Phase contrast shows the morphology of the cells stained.

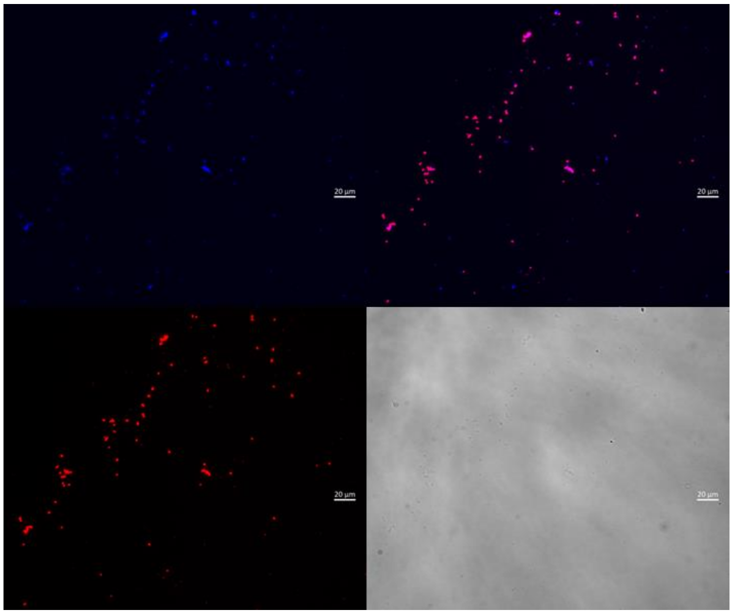

***Figure S9**. **FISH staining:** Microscopic FISH image of samples from the reactor. Blue panel represents eubacteria using a blend of EUB338, EUB338-II, and EUB338-III probes. Red panel represents “Ca. Accumulibacter” employing mixtures of the probes Acc1011, Acc471, Acc471_2, Acc635, Acc470. Magenta represents the overlap of both eubacteria and “Ca.* Accumulibacter” cells. Phase contrast shows the morphology of the cells stained.

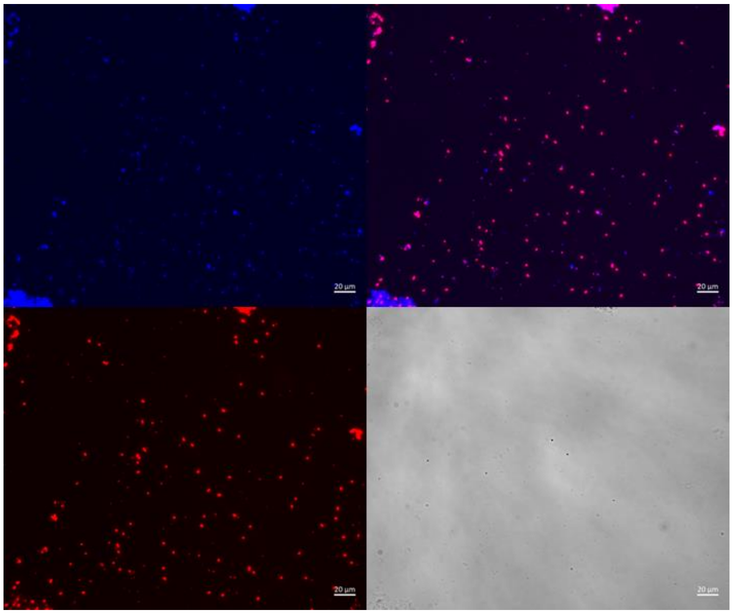

***Figure S10**. **FISH staining:** Microscopic FISH image of samples from the reactor. Blue panel represents eubacteria using a blend of EUB338, EUB338-II, and EUB338-III probes. Red panel represents “Ca. Accumulibacter” employing mixtures of the probes Acc1011, Acc471, Acc471_2, Acc635, Acc470. Magenta represents the overlap of both eubacteria and “Ca.* Accumulibacter” cells. Phase contrast shows the morphology of the cells stained.

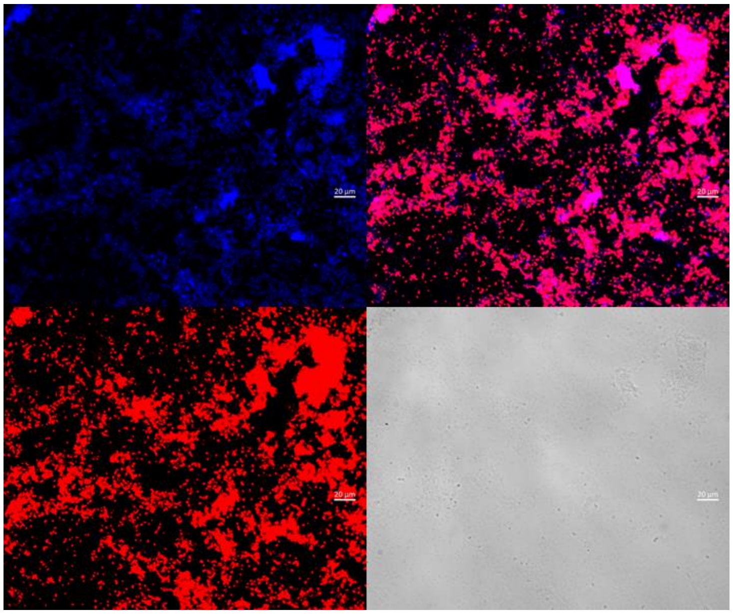

***Figure S11*. *FISH staining:*** *Microscopic FISH image of samples from the reactor. Blue panel represents eubacteria using a blend of EUB338, EUB338-II, and EUB338-III probes. Red panel represents "Ca. Accumulibacter" employing mixtures of the probes Acc1011, Acc471, Acc471_2, Acc635, Acc470. Magenta represents the overlap of both eubacteria and "Ca.* Accumulibacter" cells. Phase contrast shows the morphology of the cells stained.

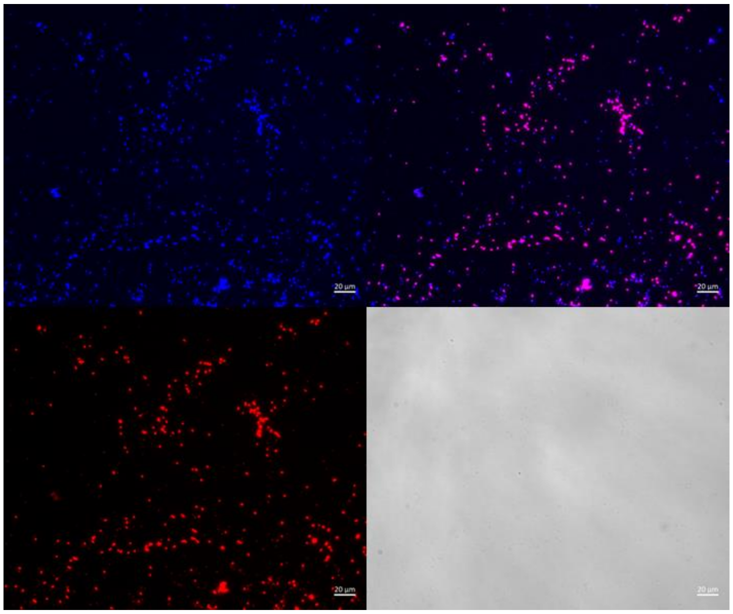

***Figure S12*. *FISH staining:*** *Microscopic FISH image of samples from the reactor. Blue panel represents eubacteria using a blend of EUB338, EUB338-II, and EUB338-III probes. Red panel represents "Ca. Accumulibacter" employing mixtures of the probes Acc1011, Acc471, Acc471_2, Acc635, Acc470. Magenta represents the overlap of both eubacteria and "Ca.* Accumulibacter" cells. Phase contrast shows the morphology of the cells stained.
